# Diurnality reconfigures circadian network dynamics in the suprachiasmatic nucleus

**DOI:** 10.64898/2026.01.28.702429

**Authors:** Vanessa Muhl, Daniela Lopez Lorenzo, Farina Pourmir, Samantha O. Sweck, Daisuke Ono, Jeff R. Jones

## Abstract

Diurnal and nocturnal mammals occupy opposite temporal niches, but whether the suprachiasmatic nucleus (SCN), the central circadian pacemaker, follows the same intrinsic rules for timing adjustment and network coordination across species is unknown. Because SCN clock gene rhythms and population-level activity measurements look broadly similar across species, the prevailing interpretation in comparative studies has been that temporal niche differences are implemented upstream or downstream of the SCN. However, these coarse readouts do not test how clock resetting depends on circadian phase or how neuronal timing is coordinated across SCN space. Here, we combined long-duration ex vivo SCN recordings with optogenetic stimulation to compare SCN network dynamics in the nocturnal mouse (*Mus musculus*) and the diurnal four-striped grass mouse (*Rhabdomys pumilio*). *Rhabdomys* SCN molecular clock rhythms exhibited a longer intrinsic period and, under the same daily stimulation protocol, converged on a larger absolute phase angle relative to the stimulus. Molecular clock phase response curves differed in overall structure across circadian time between species, including pronounced early subjective day delays in *Rhabdomys* when *Mus* responses were minimal. Phase mapping of single-cell SCN calcium rhythms revealed species-specific spatial timing organization, with a graded dorsomedial-to-ventrolateral phase progression in *Rhabdomys* and a sharper transition in *Mus*. The *Mus* and *Rhabdomys* SCN can thus appear similar by coarse readouts yet diverge in phase-dependent molecular clock resetting and network timing organization. These results indicate that models placing temporal niche divergence solely outside the SCN are mechanistically incomplete.

## INTRODUCTION

Across species, animals occupy different temporal niches (diurnal, day-active, versus nocturnal, night-active) that shift the timing of behavioral and physiological outputs such as sleep-wake and feeding [1,2]. In mammals, these near-24 h rhythms are coordinated by the suprachiasmatic nucleus (SCN), the central circadian pacemaker [3,4]. Light and other environmental cues adjust SCN network timing, and SCN outputs organize daily rhythms across the brain and body [5]. However, most mechanistic insight into SCN function comes from a small set of nocturnal rodent models, so whether diurnal mammals share the same intrinsic SCN network dynamics remains unclear.

Despite occupying opposite temporal niches, both day-active and night-active mammals can entrain to the light-dark cycle, so temporal niche preference cannot be explained by entrainment alone [6]. Instead, species differ in the circadian time-dependence and magnitude of behavioral phase shifting. For example, in many nocturnal mammals, light delivered during the subjective day produces little or no change in the phase of locomotor activity (a “dead zone”), whereas many diurnal mammals exhibit substantial subjective-day responsiveness [7–10]. These differences could reflect how photic input is gated before reaching the SCN, how SCN output is interpreted by circuits that control diurnal and nocturnal behavior, or how SCN coupling and circadian state shape when and how strongly the SCN resets in response to timing cues.

Because the SCN is often treated as broadly conserved in diurnal and nocturnal mammals, the prevailing view is that differences in temporal niche preference arise primarily outside the SCN [11]. For example, across species, many SCN clock genes and clock-controlled genes peak at similar times relative to the light-dark cycle, and neural measures such as multiunit recording, whole-cell recording, and metabolic readouts typically show higher SCN activity during the day than at night [12–19]. These observations have been used to argue that diurnality and nocturnality primarily reflect how photic and nonphotic signals are routed to the SCN and how SCN output is interpreted by downstream circuits. However, these measures are typically based on a small number of timepoints and population-averaged signals, so they cannot detect species differences in circadian time-dependent resetting and the spatiotemporal organization of neuronal activity rhythms.

Directly testing this interpretation requires continuous recordings paired with controlled perturbations and cell-level spatial phase mapping in a diurnal rodent model that can be closely compared with nocturnal laboratory mice (*Mus musculus*). The four-striped grass mouse (*Rhabdomys pumilio*) is a reliably day-active murid rodent closely related to *Mus* that has been used as a model organism for circadian neuroscience [12,15,20–23]. These studies report similar SCN organization across the two species, including overall neuroanatomy and a conserved day-night organization of population activity. In addition, AAV-mediated neuronal transduction has been demonstrated in *Rhabdomys*, enabling viral delivery of reporters and actuators for mechanistic experiments [24].

Here we compared *Mus* and *Rhabdomys* to test whether temporal niche preference is associated with species differences in SCN resetting and network-level timing organization. We first asked whether SCN molecular clock rhythms differ in free-running period and in the stable phase angle reached under an identical daily stimulation protocol. We then determined whether the SCN molecular clock phase response curve to acute stimulation differs across circadian time between species. Finally, we tested whether neuronal activity rhythms exhibit species differences in phase organization within the SCN, consistent with differences in coupling architecture or spatial coordination. Together, these experiments evaluate whether temporal niche preference can be attributed primarily to mechanisms upstream or downstream of the SCN, or whether intrinsic SCN dynamics also differ between diurnal and nocturnal species.

## RESULTS

### Coarse phase profiles of SCN activity and the molecular clock are similar across species

Comparative studies often conclude that diurnal and nocturnal mammals share similar SCN timing by measuring SCN activity markers and clock proteins at a few timepoints across the day. We therefore quantified c-FOS (an immediate early gene product used as a marker of SCN activation) and PER2 (a core clock protein) immunostaining in the SCN of the nocturnal house mouse (*Mus musculus*) and the diurnal four-striped grass mouse (*Rhabdomys pumilio*) collected at zeitgeber times (ZT; where ZT 0 is defined as lights on) 0, 6, 12, and 18 (**Fig. 1; Supplementary Fig. 1**). In both species, c-FOS labeling varied strongly with time of day and was highest during the light phase (one-way ANOVA, p < 0.0001; **Fig. 1a-d**). PER2 labeling also varied with time of day and peaked near ZT 12 in both species (one-way ANOVA, p < 0.0001; **Fig. 1e-h**). DAPI^+^ cell counts did not vary with time of day in either species (**Supplementary Fig. 1**), supporting that our observed time-of-day-dependent changes in c-FOS and PER2 labeling reflect changes in marker-positive cells rather than changes in total cell counts. Across timepoints, overall labeling magnitude differed by species, with *Mus* showing higher c-FOS labeling than *Rhabdomys* at all ZTs and higher PER2 labeling at ZTs 6, 12, and 18 (c-FOS, two-way ANOVA for species, p < 0.0001; PER2, two-way ANOVA for species, p = 0.0005; **Fig. 1b,d,f,h**). Despite these magnitude differences, the phase profile was similar in both species, with c-FOS highest during the day and PER2 highest around dusk.

**Figure 1.**
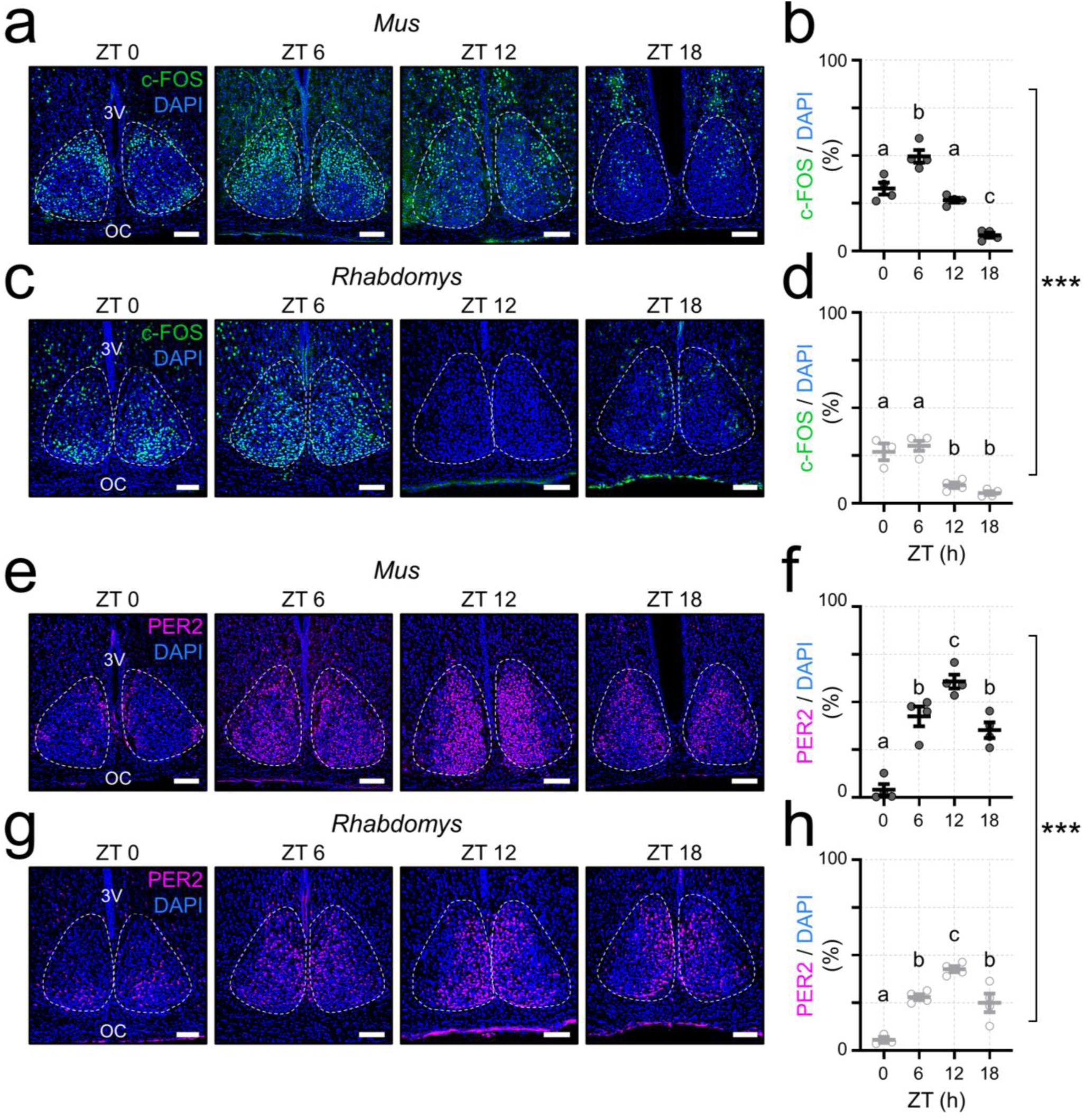
Time-of-day profiles of SCN c-FOS and PER2 labeling are aligned across species. **a,c)** Representative coronal SCN sections from *Mus* (**a**) and *Rhabdomys* (**c**) collected at ZT 0, 6, 12, and 18 under a 12 h:12 h light:dark cycle (ZT 0, lights on). c-FOS immunostaining, green; DAPI, blue. Dashed white lines outline the SCN. 3V, third ventricle; OC, optic chiasm. Scale bar, 100 µm. **b,d**) Quantification of SCN c-FOS labeling in *Mus* (**b**) and *Rhabdomys* (**d**), expressed as 100 × (c-FOS^+^ cell count / DAPI^+^ cell count) within the SCN. Each point represents one animal (*Mus*: n = 4 per timepoint; *Rhabdomys*: n = 3 at ZT 0 and n = 4 at ZT 6, 12, and 18). One-way ANOVA within each species: *Mus*, F(3, 12) = 47.74, p < 0.0001; *Rhabdomys*, F(3, 11) = 27.84, p < 0.0001. Different letters indicate timepoints that differ by Tukey’s post hoc tests (p < 0.05). **e,g)** Representative coronal SCN sections from *Mus* (**e**) and *Rhabdomys* (**g**) collected at ZT 0, 6, 12, and 18. PER2 immunostaining, magenta; DAPI, blue. Dashed white lines outline the SCN. Scale bar, 100 µm. **f,h)** Quantification of SCN PER2 labeling in *Mus* (**f**) and *Rhabdomys* (**h**), computed as 100 × (PER2^+^ cell count / DAPI^+^ cell count) within the SCN (*Mus*: n = 4 per timepoint; *Rhabdomys*: n = 3 at ZT 0 and n = 4 at ZT 6, 12, and 18). One-way ANOVA within each species: *Mus*, F(3, 12) = 34.41, p < 0.0001; *Rhabdomys*, F(3, 11) = 23.96, p < 0.0001. Different letters indicate timepoints that differ by Tukey’s post hoc tests (p < 0.05). Two-way ANOVA across species and timepoints confirmed strong time-of-day modulation for both markers and a difference in overall labeling magnitude between species (c-FOS: interaction F(3, 23) = 5.829, p = 0.0041; time F(3, 23) = 71.45, p < 0.0001; species F(1, 23) = 43.50, p < 0.0001. PER2: interaction F(3, 23) = 2.660, p = 0.0721; time F(3, 23) = 55.52, p < 0.0001; species F(1, 23) = 16.40, p = 0.0005).

### Species differences in SCN molecular clock period and phase angle of entrainment

Given this apparent similarity in static measurements of SCN timing, we next asked whether intrinsic SCN molecular clock dynamics differed across species. We prepared organotypic SCN slices from *Mus* and *Rhabdomys*, then transduced slices with AAVs encoding *Per2*-ELuc, a bioluminescent molecular clock reporter (and, where indicated, AAV-ChrimsonR, a red-shifted optogenetic activator) [12,25]. We then performed long-term bioluminescence recordings of *Per2-*ELuc rhythms (**Fig. 2a,b**). Under baseline conditions, *Rhabdomys* SCN slices exhibited a longer *Per2*-ELuc free-running period than *Mus* (**Fig. 2c-e**; Welch’s t test, p < 0.0001), whereas rhythm amplitude as measured by χ^2^ periodogram peak value did not differ between species (**Fig. 2f**; Welch’s t test, p = 0.7793).

**Figure 2.**
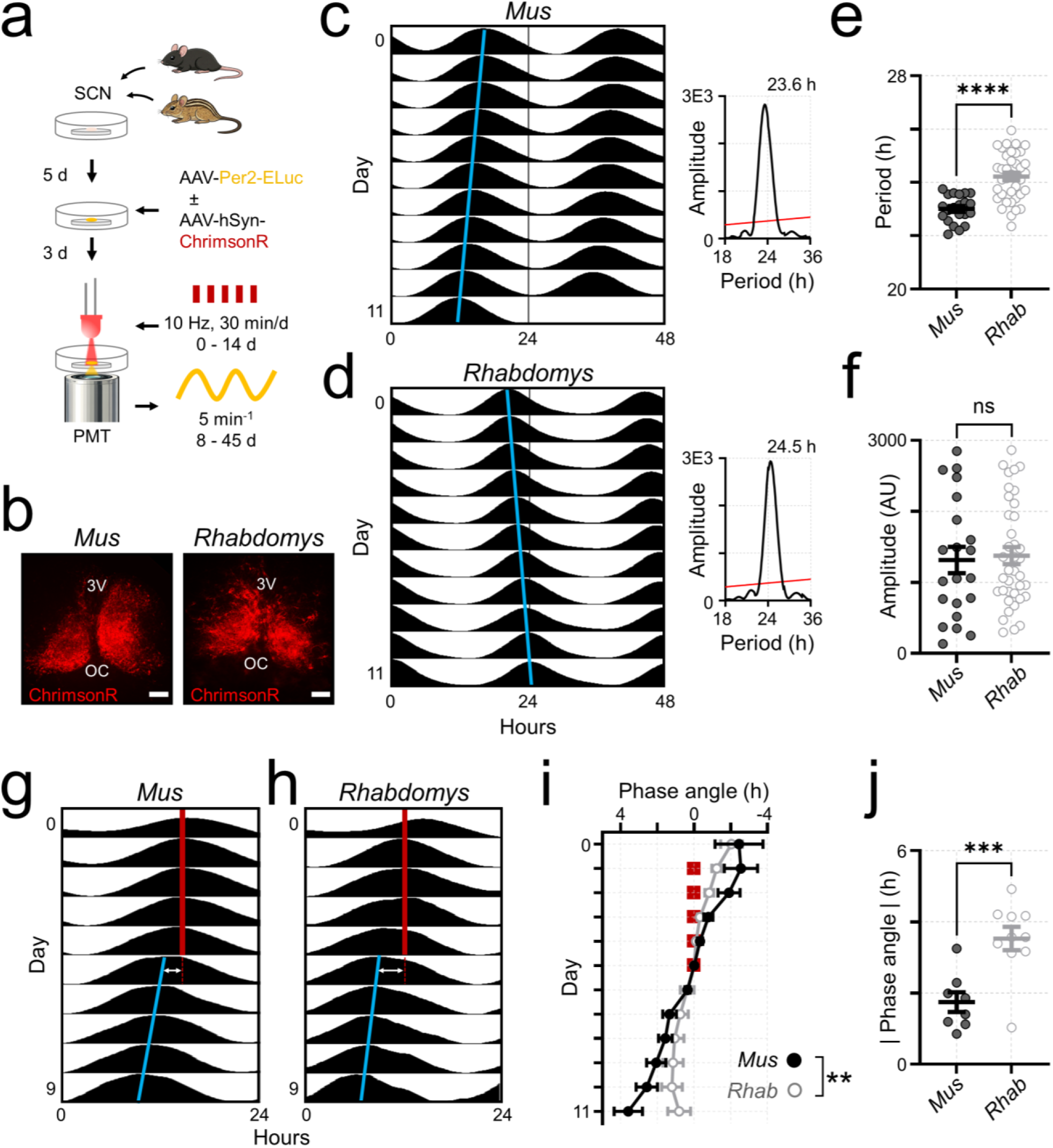
SCN molecular clock period and phase angle of optogenetic entrainment differ between nocturnal and diurnal mice. **a)** Experimental overview. Organotypic suprachiasmatic nucleus (SCN) slices were prepared from the nocturnal house mouse (*Mus musculus*) and the diurnal four-striped grass mouse (*Rhabdomys pumilio*). After 5 days in vitro, slices were transduced with AAVs encoding the bioluminescent molecular clock reporter *Per2*-ELuc alone, or together with AAV-ChrimsonR for optogenetics experiments. After an additional 3 days, slices were transferred to photomultiplier tube (PMT) recording chambers for bioluminescence acquisition (5 min bins). For optogenetics experiments, slices received daily optical stimulation (10 Hz, 30 min, 625 nm) for the indicated number of days. **b)** Representative images of ChrimsonR expression (red) in *Mus* (left) and *Rhabdomys* (right) SCN slices. Scale bar, 100 µm. 3V, third ventricle; OC, optic chiasm. **c-d)** Representative Per2-ELuc bioluminescence actograms from a *Mus* SCN slice (**c**) and a *Rhabdomys* SCN slice (**d**). Actograms are double-plotted (48 h per row). Blue lines indicate peak times, defined as the local maximum of smoothed bioluminescence. Insets depict χ^2^ periodograms with the dominant period peak (black trace) and significance threshold (red line). **e)** Free-running period of *Per2*-ELuc rhythms in SCN slices from *Mus* (n = 21 animals) and *Rhabdomys* (n = 38 animals) computed from the χ^2^ periodogram peak. Each point represents one animal. Two-tailed Welch’s t test; t = 6.662, df = 56.93, p < 0.0001. **f)** χ^2^ periodogram amplitude (peak value) of *Per2*-ELuc rhythms in *Mus* (n = 21 animals) and *Rhabdomys* (n = 38 animals). Each point represents one animal. Two-tailed Welch’s t test; t = 0.2822, df = 36.7, p = 0.7793. **g-h)** Representative single-plotted *Per2*-ELuc actograms from a *Mus* SCN slice (**g**) and a *Rhabdomys* SCN slice (**h**) during daily optogenetic stimulation (10 Hz, 30 min, 625 nm; red line) and subsequent free run. Blue lines indicate peak times (local maxima of smoothed bioluminescence) after stimulation ended. Phase angle of entrainment (white double-headed arrow) was quantified on the first post-stimulation day as the time difference between the *Per2*-ELuc peak and the extrapolated stimulation time (dashed red line). **i)** Trajectory of phase angle between the daily optogenetic stimulation time and the *Per2*-ELuc peak (mean ± SEM) during entrainment and subsequent free run in *Mus* (filled circles, black; n = 8 animals) and *Rhabdomys* (open circles, gray; n = 10 animals). Red squares at 0 h indicate stimulation days (days 1 to 5). Phase angle was computed as stimulation time minus *Per2*-ELuc peak time, where positive values indicate peaks occurring before stimulation. Mixed effects model (REML) with fixed effects of time, species, and interaction: time F(11, 159) = 21.10, p < 0.0001; species F(1, 16) = 0.3622, p = 0.5557; interaction F(11, 159) = 2.478, p = 0.0068. **j)** Absolute phase angle of entrainment (hours) on day 6 (first day after the end of optogenetic stimulation) quantified for individual animals. Absolute value is plotted to summarize stable entrainment occurring on either side of the stimulation phase. *Mus* n = 8 animals; *Rhabdomys* n = 10 animals. Two-tailed Welch’s t test; t = 4.161, df = 15.96, p = 0.0007.

Clocks that run faster or slower than 24 h would drift each day unless a daily timing cue produces a corrective phase shift [26]. Because the SCN molecular clocks of *Mus* and *Rhabdomys* differed in their intrinsic periods, we next tested whether the same daily cue results in the same timing of the *Per2*-ELuc peak relative to that cue in both species. We delivered optogenetic stimulation (10 Hz, 625 nm) to ChrimsonR-expressing SCN slices for 30 min per day for 5 days (**Fig. 2a**), then quantified the timing of the first post-stimulation *Per2*-ELuc peak relative to the scheduled stimulation time (**Fig. 2g,h**) [25,27–29]. In both species, *Per2*-ELuc rhythms successfully entrained to the stimulation and free-ran with a defined phase angle of entrainment after stimulation ended, with the day-by-day trajectory differing by species (**Fig. 2i**; mixed-effects model, time-by-species interaction, p = 0.0068). Consistent with a species difference in stabilized *Per2*-ELuc timing to the same daily optogenetic timing cue, the absolute magnitude of the phase angle of optogenetic entrainment on the first post-stimulation day was greater in *Rhabdomys* than in *Mus* (**Fig. 2j**; Welch’s t test, p = 0.0007).

Together, these results establish two species-level differences in SCN molecular clock dynamics under the same recording and stimulation protocol. First, the free-running period of the SCN *Per2*-ELuc rhythm was shorter in *Mus* and longer in *Rhabdomys*. This matches the direction of long-standing behavioral observations that diurnal species tend to show longer free-running periods in constant darkness than nocturnal species, but here assessed at the level of the SCN molecular clock rather than overt behavior [30]. Second, under an identical daily optogenetic stimulation protocol, *Mus* and *Rhabdomys* SCN *Per2*-ELuc rhythms converged on distinct phase angles relative to the stimulus, indicating species differences in the stable phase relationship produced by the same periodic input.

### Species differences in SCN molecular clock phase resetting

If the same daily timing cue causes the *Per2*-ELuc rhythm to peak at different times relative to that cue in the two species, then the cue must shift their clocks differently depending on when it is delivered [31]. We therefore quantified this phase dependence directly by constructing phase response curves (PRCs) for each species [32]. We delivered a single optogenetic pulse to ChrimsonR-transduced SCN slices from *Mus* and *Rhabdomys* at defined circadian times and quantified phase shifts from changes in the regression of the daily acrophase before versus after stimulation (minimum 3 d each; **Fig. 3a,b**) [25,27–29]. Stimulation time was expressed on a fixed 24 h circadian time scale, with CT 12 anchored to the extrapolated *Per2*-ELuc peak on the day of stimulation.

**Figure 3.**
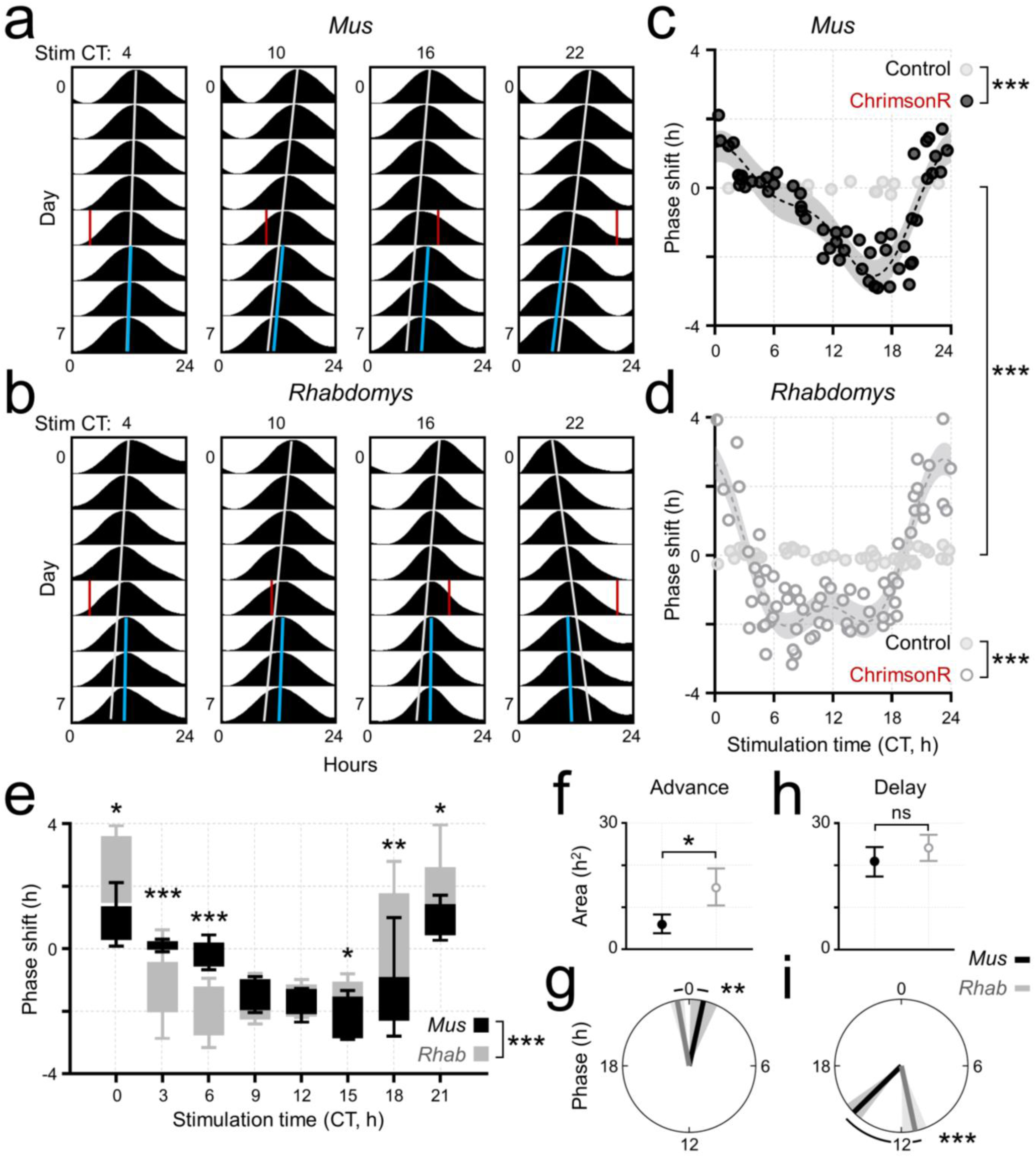
Optogenetic phase resetting of SCN molecular clock rhythms differs between nocturnal and diurnal mice. **a-b)** Representative single-plotted *Per2*-ELuc bioluminescence actograms from ChrimsonR-transduced SCN slices from nocturnal *Mus musculus* (**a**) and diurnal *Rhabdomys pumilio* (**b**) showing responses to a single optogenetic stimulation (30 min, 10 Hz, 625 nm) delivered on day 4 at the indicated circadian times (from left to right, CT 4, 10, 16, 22). Gray lines indicate the pre-stimulation rhythm extrapolated through the stimulation day, and blue lines indicate post-stimulation peak times. Peak times were defined as the local maximum of smoothed bioluminescence. CT was mapped onto a 24 h scale, with CT 12 anchored to the extrapolated *Per2*-ELuc peak on the stimulation day, independent of free-running period. **c-d)** Phase response curves (PRCs) for optogenetic stimulation of ChrimsonR-transduced SCN slices from *Mus* (**c**) and *Rhabdomys* (**d**). Points depict individual phase shifts as a function of stimulus CT (phase delays negative, phase advances positive). Phase shifts were quantified by comparing linear regressions of daily peak-time (acrophase) estimates over a minimum of 3 d pre-stimulation and 3 d post-stimulation. ChrimsonR-transduced slices are depicted as black filled circles (*Mus*) or gray open circles (*Rhabdomys*). ChrimsonR-negative control slices, which received identical light delivery and optical power, are depicted as light gray filled circles. Shaded bands indicate the 95% confidence interval of the fitted PRC (periodic Fourier regression, k = 2). *Mus*: ChrimsonR n = 54 stimulations from 26 animals, control n = 14 stimulations from 6 animals. *Rhabdomys*: ChrimsonR n = 67 stimulations from 36 animals, control n = 38 stimulations from 17 animals. PRC shape differed between ChrimsonR and control slices within each species (animal-blocked permutation test on curve distance): *Mus* ChrimsonR vs. control D_obs_ = 42.702, p = 0.0005; *Rhabdomys* ChrimsonR vs. control D_obs_ = 70.751, p = 0.0005. PRC shape also differed between species (animal-blocked permutation test on curve distance): *Mus* vs. *Rhabdomys* D_obs_ = 34.377, p = 0.0005. For animals receiving multiple single pulses, stimulations were separated by at least 4 d and CT was reestimated prior to each pulse. **e)** Binned PRCs for ChrimsonR-transduced slices. Phase shifts were grouped into 3 h CT bins and summarized as Tukey box-and-whisker plots (minimum, Q1, median, Q3, maximum) for *Mus* and *Rhabdomys*. Species differences across CT were tested by a mixed-effects model (REML) with fixed effects of CT bin, species, and interaction and a random intercept for animal: CT bin F(3.821, 57.32) = 34.18, p < 0.0001; species F(1, 105) = 2.212, p = 0.1399; interaction F(3.821, 57.32) = 8.565, p < 0.0001. Asterisks indicate Tukey post-hoc comparisons between species within a bin (* p < 0.05; ** p < 0.01; *** p < 0.001). **f-i)** Summary features extracted from the fitted mean PRCs. Uncertainty was assessed by animal-level bootstrap; between-species features were tested by animal-blocked permutation with Holm correction. **f)** Advance-zone area, computed as the integral of the positive values of the fitted PRC over CT (in h^2^) differed between species (Holm-adjusted p = 0.0240). **g)** Advance-zone centroid phase (CT), shown on a circular plot, differed between species (Holm-adjusted p = 0.0060). Arrows indicate the circular mean for each species and shaded sectors indicate the 95% bootstrap shortest-arc interval. **h)** Delay-zone area, computed as the integral of the negative values of the fitted PRC over CT (in h^2^), did not differ between species (Holm-adjusted p = 0.4008). **i)** Delay-zone centroid phase (CT), shown on a circular plot as in **g**, differed between species (Holm-adjusted p = 0.0040).

Optogenetic PRCs derived from ChrimsonR-expressing slices differed from light-only controls in both species (**Fig. 3c,d**; animal-blocked permutation test, p = 0.0005), confirming that phase resetting depended on optogenetic activation. Surprisingly, PRC shape also differed between *Mus* and *Rhabdomys* (animal-blocked permutation test, p = 0.0005), indicating that the phase dependence of molecular clock resetting differed by species. Consistent with this, binned PRCs (3 h bins) showed a significant CT bin by species interaction (**Fig. 3e**; mixed-effects model, interaction p < 0.0001). In late subjective night and early subjective day bins (CTs 18 to 0), *Rhabdomys* exhibited larger phase advances than *Mus*. In early subjective day bins (CTs 3 and 6), *Rhabdomys* exhibited phase delays while *Mus* responses were near zero (Tukey post-hoc comparisons within bin, as indicated within **Fig. 3e**). This suggested that there may be a species difference in phase resetting during the circadian “dead zone,” a time interval where light typically produces little or no phase shift in *Mus* behavioral rhythms [33].

To test this directly, we defined *a priori* an early subjective day window (CT 2 to 8) classically considered the circadian dead zone [32]. Importantly, we use dead zone here as an operational label for this prespecified CT interval, not as a claim about the absence of resetting under all stimuli. Within this window, *Rhabdomys* exhibited stronger delays than *Mus* (mean delay magnitude: *Mus*, 0.16 h, *Rhabdomys*, 1.11 h; animal-blocked permutation test, p = 0.0064). This conclusion was supported by other prespecified window metrics (mean signed shift, p = 0.0108; mean delay area, p = 0.0064).

Next, to quantify the timing and size of the advance and delay zones, we extracted features from the fitted PRCs (**Fig. 3f-i**). The advance-zone area was greater in *Rhabdomys* than in *Mus* (animal-blocked permutation test; p = 0.0240; **Fig. 3f**), and the phase of the advance-zone centroid (the center of mass of the advance-zone) was earlier in *Rhabdomys* (CT 23.32 vs. 0.82; **Fig. 3g**; animal-blocked permutation permutation test, p = 0.0060). The delay-zone area did not differ between species (**Fig. 3h**; animal-blocked permutation test, p = 0.4008), but the phase of the delay-zone centroid was earlier in *Rhabdomys* (CT 11.18 vs. 15.05; **Fig. 3i**; animal-blocked permutation test, p = 0.0020).

Together, these results indicate that *Mus* and *Rhabdomys* differ in both when and how strongly the SCN molecular clock advances or delays in response to the same optogenetic stimulus. This species difference in PRC shape was driven mainly by the circadian timing of advances and delays, and by larger advances in *Rhabdomys*, rather than by a uniform rescaling of the entire PRC. Notably, *Rhabdomys* exhibited stronger delays than *Mus* in the early subjective day, consistent with a between-species difference in resetting during an interval classically associated with minimal light-induced behavioral phase shifts in nocturnal rodents.

### Species differences in phase resetting are not driven by period shifts

Because resetting stimuli can also modulate period, we asked whether stimulation-evoked changes in free-running period could account for the species differences in PRC shape [34]. We constructed period response curves (τRCs) using the same stimulation dataset and the same regression-based estimator used for phase-shift quantification in **Fig. 3**. Within each species, τRC shape differed between ChrimsonR-expressing slices and light-only controls (**Supplementary Fig. 2a,b**; animal-blocked permutation test, p = 0.0005), confirming that period modulation depended on optogenetic activation. In contrast, τRC shape did not differ between species (animal-blocked permutation test, p = 0.0970), arguing against a species-specific period effect as the primary driver of our observed PRC differences.

This interpretation is supported by three additional observations. First, period shifts covaried with phase shifts in both species with indistinguishable slopes (**Supplementary Fig. 2c,d**; ANCOVA interaction, p = 0.9984). Second, phase shifts covaried with the initial pre-stimulation period within each species, with no species difference in slope (**Supplementary Fig. 2e,f**; ANCOVA interaction, p = 0.2730). Third, the initial pre-stimulation period (z-scored within species to account for baseline period differences) did not differ between *Mus* and *Rhabdomys* in its dependence on stimulation time (**Supplementary Fig. 2g,h**; animal-blocked permutation test, p = 0.1499). Together, these analyses indicate that species differences in optogenetic phase resetting are unlikely to be explained by species-specific period modulation.

### Spatial organization of SCN calcium rhythm phase differs by species

Phase resetting reflects bidirectional coupling between SCN firing rate rhythms and the molecular clock: changes in neuronal activity can shift molecular clock phase, and the molecular clock in turn helps organize the daily pattern of SCN activity [19,27]. Because we observed species differences in molecular clock resetting, we next asked whether the spatial phase organization of single-neuron calcium rhythms, a readout of rhythmic SCN activity, differs between *Mus* and *Rhabdomys* [35–37]. To test this, we performed long-term single-cell calcium imaging of GCaMP6s-transduced SCN slices and observed robust rhythmic activity across the SCN in both species (**Fig. 4a,b**). In representative SCN slices, when single-neuron rhythms were ordered from dorsomedial to ventrolateral within each slice, peak timing appeared to progress across the tissue, with qualitatively different phase profiles in *Mus* and *Rhabdomys* (**Fig. 4c,d**). We therefore performed a quantitative, atlas-registered phase-mapping analysis to test whether spatial phase organization differs between species at the group level (**Fig. 4e**; workflow summarized in **Supplementary Fig. 3**).

**Figure 4.**
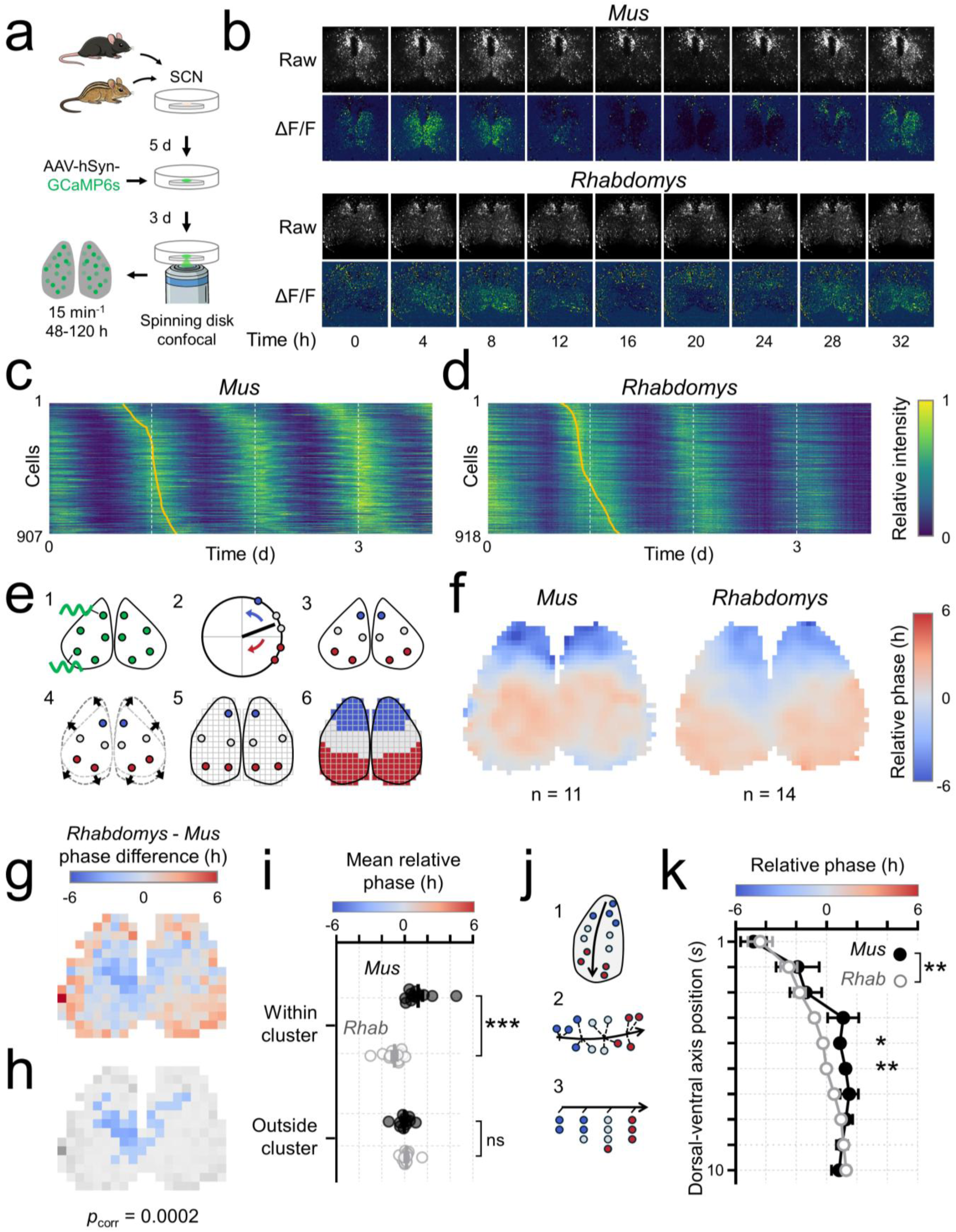
SCN single-neuron calcium rhythm phase organization differs between nocturnal and diurnal mice. **a)** Experimental overview. Organotypic SCN slices from nocturnal *Mus musculus* and diurnal *Rhabdomys pumilio* were transduced with AAVs encoding the genetically-encoded calcium indicator GCaMP6s and imaged at single-cell resolution (one frame every 15 min for 48 to 120 h). **b)** Representative GCaMP6s images from *Mus* (top) and *Rhabdomys* (bottom) SCN slices across time ex vivo (0 to 32 h in 4 h increments). For each time point, the top row shows raw fluorescence and the bottom row shows ΔF/F (normalized change in fluorescence). **c-d)** Single-cell calcium activity heatmaps from representative *Mus* (**c**) and *Rhabdomys* (**d**) SCN slices. Rows correspond to tracked neuronal ROIs (ordered dorsomedial to ventrolateral), columns correspond to recording time in days. Heatmap values depict within-cell normalized fluorescence (0 to 1) across the full recording window. Yellow line indicates the time of maximal fluorescence for each ROI on the first recording day containing a complete cycle. **e)** Workflow for phase mapping and group comparison. Rhythmic ROIs were identified using single-component Cosinor fitting (p < 0.05, r^2^ ≥ 0.2), and peak phase was defined as the Cosinor acrophase (in hours, wrapped within one cycle). Within each slice, phases were defined in units of the estimated period, then centered by subtracting the circular mean phase across the SCN to yield relative phase. ROIs were warped into a common SCN atlas coordinate frame to compare spatial phase patterns across animals. For group analysis, phases were converted to circadian time, and all binning, circular means, and phase-difference calculations were performed using a fixed 24 h cycle. **f)** Group-averaged relative phase maps in SCN atlas space for *Mus* (n = 11 animals) and *Rhabdomys* (n = 14 animals). ROIs were binned on a 40 x 40 grid; each bin depicts the circular mean of ROI-level relative phases, pooled across animals. A Gaussian spatial smoothing kernel was applied for visualization. **g)** Binwise group phase-difference map (*Rhabdomys* minus *Mus*) calculated from animal-level circular means per bin on a 20 x 20 grid. Bins required ≥5 animals per species for visualization (**g**) and ≥9 animals per species for inference (**h**), with ≥10 cells per bin. **h)** Cluster-based permutation inference on the 20 x 20 phase-difference map. The per-bin test statistic was the absolute circular phase difference. Cluster-based permutation test (10,000 permutations) controlled familywise error by max cluster mass (cluster-forming α= 0.2). The significant cluster is shown in color (cluster-corrected p = 0.00024). **i)** Per-animal circular mean relative phases within and outside the significant cluster in (**h**). Region and species effects were tested using a Harrison-Kanji two-way circular ANOVA: region F = 14.563, p = 0.0004; species F = 1.113, p = 0.2970; interaction F = 20.950, p < 0.0001. Post-hoc Watson-Williams tests comparing *Mus* vs. *Rhabdomys* identified a species difference within, (F = 23.115, p < 0.001), but not outside (F = 0.7941, p = 0.3821) the cluster. **j)** Schematic of phase-axis gradient analysis. In atlas space, a dorsomedial-to-ventrolateral phase axis was defined as a polyline. Warped ROI centroids were projected onto the nearest point on the polyline to obtain a normalized arc-length coordinate *s*, with bilateral assignment to the nearest axis and automatic orientation standardization. **k)** Phase gradient along the dorsomedial-to-ventrolateral axis. Relative phase (hours) is plotted across 10 equal-length segments along the normalized axis coordinate *s* (1 dorsomedial, 10 ventrolateral); points (*Mus*, black filled circles; *Rhabdomys*, gray open circles) depict mean ± SEM across animals. Two-way repeated measures ANOVA with Geisser-Greenhouse correction: axis position F(2.85, 65.57) = 21.56, p < 0.0001; species F(1, 23) = 7.188, p = 0.013; interaction F(2.85, 65.57) = 0.882, p = 0.4506. Sidak post hoc tests identified species differences at positions 5 (p = 0.016) and 6 (p = 0.002).

Group-averaged atlas-space phase maps of single-cell calcium rhythms suggested that *Mus* and *Rhadomys* differ in how relative phase (phase relative to the slice mean) is organized across the SCN (**Fig. 4f**; individual-animal binned maps underlying the group averages are shown in **Supplementary Fig. 4**). In *Mus*, relative phase showed a sharper transition from an advanced dorsomedial region to a delayed ventrolateral region, whereas in *Rhabdomys*, the transition appeared more gradual across the tissue. To test this quantitatively, we computed an atlas-registered phase-difference map from animal-level per-bin circular means and applied cluster-based permutation inference to identify spatially contiguous phase differences while controlling familywise error (**Fig. 4g,h**). This analysis revealed a significant cluster localized to an intermediate atlas region (cluster-corrected p = 0.00024), consistent with a spatially constrained species difference rather than a uniform phase offset. Per-animal phase summaries revealed a strong region-by-species interaction (**Fig. 4i**; Harrison-Kanji test, interaction p < 0.0001), with *Rhabdomys* phase-advanced relative to *Mus* within the cluster (Watson-Williams test, p < 0.001; mean difference ∼2.1 h) but not outside it (p = 0.3821).

Because the atlas maps suggested a dorsomedial-to-ventrolateral difference in relative phase organization between species, we next tested this pattern explicitly by quantifying relative phase along a defined anatomical axis (**Fig. 4j**). This analysis matched the atlas maps: *Rhabdomys* showed a graded dorsomedial-to-ventrolateral phase progression, whereas *Mus* showed a sharper transition (**Fig. 4k**). Accordingly, relative phase varied strongly with axis position (repeated-measures ANOVA, position p < 0.0001) and differed overall between species (species p = 0.013). This between-species difference was driven by a phase advance in *Rhabdomys* relative to *Mus* at intermediate axis positions (Sidak’s post hoc tests, positions 5 and 6, p = 0.016 and p = 0.002), consistent with the intermediate-region phase divergence identified in the 2D phase difference analysis (**Fig. 4h,i**).

Together, these results demonstrate that *Mus* and *Rhabdomys* differ in how the timing of the daily calcium peak is patterned across SCN space. In *Mus*, relative phase is organized as an earlier dorsomedial region and a later ventrolateral region with a comparatively abrupt transition, consistent with prior reports of spatial phase organization in the mouse SCN [35,37,38]. In contrast, *Rhabdomys* peak timing shifts progressively across the same dorsomedial-to-ventrolateral axis. The largest between-species divergence is concentrated in an intermediate SCN region, where *Rhabdomys* is phase-advanced relative to *Mus*.

## DISCUSSION

In this study, we used isolated ex vivo SCN explants from the nocturnal laboratory mouse *Mus musculus* and the diurnal four-striped grass mouse *Rhabdomys pumilio* to test whether intrinsic properties of the central circadian pacemaker are conserved across species occupying different temporal niches. In preparations that exclude retinal input and disconnect the SCN from its normal downstream targets, we observed species differences in SCN resetting dynamics and network-level timing organization. These findings constrain prevailing interpretations based on coarse SCN measures, such as timepoint immunostaining of SCN activation and clock protein expression, and demonstrate that broad similarity between species in SCN timing does not imply conserved dynamical rules for when and how strongly timing cues can shift the clock, nor conserved spatial organization of neuronal timing within the SCN.

One of the clearest species-level differences we observed was the intrinsic free-running period of the SCN molecular clock, where *Rhabdomys Per2*-ELuc rhythms had a longer free-running period than *Mus*. This direction matches Aschoff’s long-standing behavioral generalization that diurnal animals often free run in constant darkness with periods modestly longer than 24 h, whereas nocturnal animals often free run with periods modestly shorter than 24 h [30]. Although this pattern is well-established at the behavioral level, direct tests at the level of the SCN molecular clock in diurnal mammals have been limited, in part because many cross-species studies rely on sparse time-point measurements of clock gene expression that resolve relative phase but cannot estimate free-running period with high precision. Our long-duration *Per2*-ELuc recordings therefore show that this difference in free-running period can be expressed by the SCN molecular clock itself.

We also found that daily stimulation could entrain the SCN molecular clock rhythm in both *Mus* and *Rhabdomys*, consistent with prior demonstrations of optogenetic entrainment in the ex vivo *Mus* SCN [25]. After several days, *Per2*-ELuc peaks converged on a stable phase angle relative to the scheduled stimulation. This offset differed by species, with *Rhabdomys* exhibiting a larger absolute phase angle than *Mus*. A stable phase angle is expected when the clock’s intrinsic tendency to drift each day (because its period is not exactly 24 h) is balanced by the timing adjustment produced by the daily stimulation [39]. Because that adjustment depends on the oscillator’s internal circadian state, species differences in intrinsic period and state-dependent responsiveness plausibly contribute to the species-specific phase angles reached under an identical stimulation protocol.

To test state dependence directly, we constructed phase response curves (PRCs) that quantify how the same SCN stimulus shifts molecular clock timing as a function of circadian time. In *Mus*, the *Per2*-ELuc PRC evoked by optogenetic stimulation closely resembles PRCs reported in prior ex vivo optogenetic-resetting studies using SCN from transgenic PER2::LUC mice, supporting that our AAV-based reporter reproduces canonical SCN molecular clock resetting dynamics [25,27–29]. In *Rhabdomys*, however, the PRC differed from *Mus* in overall structure, not through a uniform change in phase-shift magnitude or a rigid translation along the circadian time axis. The advance zone was larger, the delay zone was similar in overall size, and both zones occurred earlier in circadian time, but were shifted by different amounts. This pattern indicates a species difference in how circadian state sets the sensitivity of the SCN molecular clock to a standardized perturbation delivered to the network.

A particularly informative consequence of the species difference in PRC structure is that the ability to reset the clock differs during the early subjective day, an interval often treated as a “dead zone” for light-induced behavioral phase shifts in nocturnal rodents. The same optogenetic stimulation produced little to no shift in *Mus* SCN *Per2*-ELuc rhythms, but produced a pronounced delay in *Rhabdomys* rhythms, indicating that this time window permits robust SCN molecular clock resetting in *Rhabdomys* but not in *Mus*. Because the stimulus was delivered directly to isolated SCN tissue, this indicates that the diurnal versus nocturnal difference in subjective-day responsiveness can be generated within the SCN network itself without requiring retinal phototransduction or downstream decoding of SCN outputs.

Our single-cell timing maps show that *Mus* and *Rhabdomys* differ in how the relative phase of the SCN calcium rhythm is organized across SCN space. The divergence is regionally concentrated rather than uniformly distributed, with the largest between-species phase difference confined to an intermediate band along the dorsomedial-to-ventrolateral axis. The direction of this effect is a relative phase advance in *Rhabdomys* compared to *Mus*, consistent with the broader spatial phase profiles in each species: a graded dorsomedial-to-ventrolateral phase progression in *Rhabdomys* versus a sharper transition in *Mus*. The dorsomedial-advanced, ventrolateral-delayed topology in *Mus* aligns with prior reports of SCN calcium rhythm organization in this species [35,37,38].

Several circuit-level differences between *Mus* and *Rhabdomys* could generate these spatial phase profiles. Prior work in the *Mus* SCN shows that calcium rhythms can retain structured spatial topology even when population-scale electrical activity appears comparatively synchronous, meaning that calcium timing can reveal network organization not captured by coarse electrical readouts that can be broadly aligned across species [37]. Accordingly, a phase-advanced intermediate region in *Rhabdomys* is compatible with differences in coupling strength or coupling topology across SCN compartments, differences in how oscillator subpopulations set local phase, or spatial heterogeneity in intrinsic oscillator properties that stabilizes timing differently across the tissue [35,40–42]. This is consistent with our discovery of species differences in molecular clock resetting because local perturbations become tissue-level phase shifts only after being integrated across coupled SCN regions, and that process is constrained by spatial phase organization and network coupling.

Species-level conclusions about resetting and spatial timing require inference that preserves circadian phase, spatial structure, and the animal as the unit of replication. Accordingly, we treated PRCs as continuous periodic functions (rather than bin-wise summaries), and tested species divergence with animal-blocked resampling [43,44]. We then summarized PRC structure with a small set of prespecified, interpretable features under explicit multiplicity control to prevent overinterpretation of feature-level effects. For calcium timing, we estimated phase at the level of single neuronal ROIs, registered those estimates into a shared SCN atlas, and evaluated species differences using animal-level summaries. This choice is important because slice-specific pixel gradients, uneven sampling, and other idiosyncrasies can potentially produce misleading spatiotemporal patterning, but atlas-based animal-level inference asks whether the spatial pattern reproduces across animals in a common anatomical framework.

One limitation of our study is that while we included males and females of each species, our experiments were not designed or powered to test sex as a modifier of SCN resetting or spatial timing organization. Sex effects, including potential sex-by-species interactions in these endpoints, therefore remain unresolved. In addition, ex vivo SCN slices isolate intrinsic network dynamics by design, but they remove systemic inputs and feedback that are present in vivo. As such, these data do not on their own establish how our observed species differences translate into behavioral phase shifting or temporal niche preference without direct in vivo measurements or manipulations that link SCN dynamics to behavior [45]. Finally, spatial inference depends on atlas registration and on analysis choices that trade spatial resolution for statistical robustness. Our animal-level calcium rhythm maps and coverage constraints emphasize reproducible, group-level organization, but they may be less sensitive to finer-grained, cell-type specific spatial structure.

Prior comparative work has often interpreted temporal niche differences as arising downstream from the SCN because diurnal and nocturnal mammals show broadly similar SCN properties in coarse readouts, including clock gene rhythm timing relative to the light-dark cycle, population activity rhythms that are high during the day, and gross neuroanatomy. However, these measures lack the temporal continuity and spatial resolution needed to test whether species differ in circadian time-dependent resetting of the molecular clock or in the rules that coordinate timing across SCN space. Here we show that the *Mus* and *Rhabdomys* SCN can appear similar by these readouts yet diverge in phase-dependent optogenetic resetting of *Per2-*ELuc rhythms and in the spatial organization of single-cell calcium rhythm timing. We do not argue that the SCN alone determines diurnality or nocturnality, but our data demonstrate that intrinsic SCN network dynamics and coordination differ between species in isolated slices under a matched recording and stimulation protocol. Models that place temporal niche preference solely upstream or downstream of the SCN are therefore mechanistically incomplete.

## METHODS

### Animals

The original four-striped grass mouse (*Rhabdomys pumilio*) breeding pairs used to establish our colony were provided by the Mallarino lab (Princeton University) [46]. The Princeton colony originated from F10 descendants of wild-derived animals originating from Goegap Nature Reserve, South Africa (S 29° 41.56′, E 18° 1.60′). Adult *Rhabdomys* and laboratory mice (wild-type *Mus musculus*; on a C57BL6/J background) were housed under a 12 h:12 h light:dark cycle (LD; lights on defined as zeitgeber time 0) at constant temperature (∼23°C) and humidity (∼40%) with food and water provided *ad libitum*. Light intensity measured at the cage floor was ∼3 x 10^14^ photons/cm^2^/s. Male and female pups of both species (postnatal days 14 to 21) were used for organotypic slice cultures and subsequent experiments. All experiments were approved by and performed in accordance with the guidelines of Texas A&M University’s Institutional Animal Care and Use Committee.

### Timepoint perfusion, immunohistochemistry, and imaging

Adult *Mus musculus* and *Rhabdomys pumilio* (2-7 months old; *Mus*, n = 16; *Rhabdomys*, n = 15) were housed in custom-built, light-tight circadian cabinets with controlled temperature and humidity under either a 12 h:12 h light:dark schedule (lights on 07:00, ZT 0) or a reverse 12 h:12 h schedule (lights on 19:00, ZT 0) for at least 7 d before perfusion. Animals were euthanized by isoflurane overdose at ZT 0, 6, 12, or 18; ZT 12 and ZT 18 procedures were performed under dim red light (650 nm). Animals were transcardially perfused with phosphate-buffered saline (PBS) followed by 4% paraformaldehyde (PFA). Brains were extracted, post-fixed overnight in 4% PFA, cryoprotected in 30% sucrose, and sectioned coronally at 40 μm on a cryostat. Free-floating sections were processed for immunostaining using established methods [47]. Primary antibodies were rabbit anti-PER2 (1:1000; AB2202, Millipore) and guinea pig anti-c-FOS (1:1000; 226 005, Synaptic Systems). Secondary antibodies were CF568 goat anti-rabbit (1:500; 20103, Biotium) and CF488A goat anti-guinea pig (1:500; 20489, Biotium). All brains were processed in parallel using identical antibody concentrations, incubation durations, and wash conditions. Sections were imaged on a Nikon Ti2-E equipped with an X-Light V3 spinning-disk confocal (Crest Optics), using acquisition settings held constant across animals and timepoints.

### Image quantification

For each animal, we quantified labeling in the mid-SCN from 5 to 8 SCN-containing coronal sections (section counts varied because occasional tissue loss occurred during sectioning or immunohistochemistry). For each section, the SCN was outlined in Fiji (ImageJ) and analyzed with a fixed, channel-specific workflow applied identically to all images. Within each channel, images were background-subtracted and binarized using automated thresholding. Objects were then quantified using the Analyze Particles function using prespecified size and circularity constraints to count c-FOS or PER2-positive cells or DAPI-stained nuclei while excluding debris. For each marker, left and right SCN counts were combined within each section and then averaged across sections to yield a single per-animal estimate. Each animal contributed one value to group analysis.

### Organotypic SCN slice preparation and viral transduction

Organotypic brain slices were prepared as previously described for *Mus* [47,48]. Briefly, brains from *Mus* and *Rhabdomys* pups were rapidly removed and blocked to isolate the hypothalamus in ice-cold HBSS supplemented with 100 U/mL penicillin-streptomycin, 5 mM HEPES, and 4.5 mM sodium bicarbonate. Coronal hypothalamic slices (200 to 250 μm) containing the SCN were cut on a vibroslicer (Campden Instruments) in cold HBSS (4 to 8°C), trimmed to ∼1.5 mm^2^, and transferred to PTFE culture membrane inserts (Millipore) in 35 mm dishes containing DMEM (D5030, Sigma) supplemented with 25 U/mL penicillin-streptomycin, 10 mM HEPES, 2% B27 Plus (Gibco), 4 mM GlutaMAX (Gibco), and 4.2 mM sodium bicarbonate. For bioluminescence recordings, culture medium was exchanged immediately before the start of recording for identical medium additionally supplemented with D-luciferin (0.1 mM, GoldBio). Dishes were sealed with optically clear PCR plate film (Bio-Rad) and maintained in a non-humidified incubator at 36°C.

Slices were virally transduced by briefly removing the plate film and pipetting 1 μL of AAV (viruses detailed below) directly onto the SCN, then returning the dishes to the incubator to allow for viral expression. For *Per2*-ELuc-only or GCaMP experiments, slices received AAVs encoding *Per2*-ELuc or GCaMP6s, were returned to the incubator for 3 to 7 days, then were used for bioluminescence recording or calcium imaging. For optogenetic stimulation experiments, slices first received AAV encoding ChrimsonR and were returned to the incubator for 3 to 7 days. Slices then received AAV encoding *Per2*-ELuc and were transferred to a 32-channel, 4-photomultiplier tube luminometer (LumiCycle; Actimetrics) maintained at 36°C for an additional 3 to 7 days to allow for reporter expression, after which baseline bioluminescence recording was collected followed by optogenetic stimulation.

GCaMP6s was delivered using an hSyn1-driven GCaMP6s construct (hSyn1-GCaMP6s-2A-NLS-dTomato-WPRE-hGH-pA; Addgene #51084) packaged in AAV-PHP.eB (titer 1.80 × 10^13^ vg/mL). *Per2*-ELuc was delivered using mPer2-ELuc-PEST-SV40-pA (construct provided by Daisuke Ono, Nagoya University) packaged in AAV-PHP.eB (titer 6.10 × 10^12^ vg/mL). ChrimsonR was delivered using an hSyn1-driven ChrimsonR-tdTomato construct (hSyn1-ChrimsonR-tdTomato-WPRE-SV40-pA; derived from Addgene #92207) packaged in AAV1 (titer 6.40 × 10^12^ vg/mL). Reporter fluorophores (dTomato or tdTomato) were used to confirm transduction in culture. All AAVs were packaged by the University of Zurich Viral Vector Facility.

### Per2-ELuc bioluminescence recording and optogenetic stimulation

After viral expression of *Per2*-ELuc, bioluminescence was recorded from individual SCN slice cultures using a LumiCycle luminometer (Actimetrics) maintained at 36°C in a non-humidified, light-tight incubator according to established methods [27]. Photon counts were acquired in 5-min bins with 1-min integration (12 samples per hour). Continuous recordings lasted 8 to 45 days, and all optogenetics experiments included at least 3 baseline days of recording before stimulation.

Optogenetic stimulation was delivered using an integrated LumiCycle add-on (AM2-LCOP, Actimetrics) that selectively illuminated designated culture dishes, while the system remained light-tight outside scheduled stimulation. Red light (625 nm) was delivered at 10 Hz (10 ms pulse width) for 30 min per stimulation bout, centered over the culture dish (6 mm illumination radius). Irradiance at the level of the slice was 1.5 mW/mm^2^, measured with a PM100D power meter (Thorlabs). Dishes remained sealed with optically clear PCR plate film during stimulation. Slice surface temperature measured immediately before and after stimulation did not change within measurement resolution (± 0.1°C).

Circadian time (CT) of stimulation was assigned *post hoc* from the pre-stimulation bioluminescence rhythm (see below) because stimulation was scheduled by clock time rather than by real-time CT estimates. This approach yielded stimulation bouts spanning a range of CTs across slices. For phase response curve experiments, each slice received a single stimulation bout on one day, and bioluminescence was recorded for at least 3 days before and after stimulation to estimate pre- and post-stimulation acrophase. For experiments in which a slice received multiple stimulation bouts, bouts were separated by at least 4 days, and CT was re-estimated from the pre-stimulation rhythm prior to each bout. Stimulation parameters were chosen based on prior SCN slice optogenetic resetting and entrainment work in *Mus* [25,27].

### Per2-ELuc rhythm processing, phase and period estimation, and PRC modeling

Raw *Per2*-ELuc bioluminescence time series were processed following established methods [25]. Briefly, time series were preprocessed in LumiCycle Analysis (Actimetrics) by baseline subtraction using a 24 h running average and smoothing using a 2 h moving average. Processed traces were exported to ClockLab (Actimetrics) for visualization and peak-based timing analyses. For visualization, baseline-subtracted rhythms were displayed as single- or double-plotted actograms. Daily acrophase was defined as the local maximum of the processed bioluminescence trace. Events were excluded if daily acrophases could not be unambiguously identified on all days required for the pre- and post-stimulation regression windows. For slices receiving multiple stimulation bouts, each bout was analyzed as an independent event with its own pre- and post-stimulation windows, and bouts were separated by at least 4 days.

Circadian time (CT) was defined on a fixed 24 h scale, with CT 12 anchored to the extrapolated *Per2*-ELuc peak on the day of stimulation, using the pre-stimulation rhythm to assign CT *post hoc*. Phase shifts evoked by optogenetic stimulation were quantified using a regression-based estimator applied to daily acrophase estimates. Linear regressions were fit to peak times over a minimum of 3 cycles before stimulation and three cycles after stimulation, and the phase shift was defined as the difference between the first post-stimulation acrophase estimated from the post-stimulation regression and the acrophase predicted for that same cycle by extrapolating the pre-stimulation regression. Phase shifts are reported using the standard PRC convention: positive values indicate phase advances (earlier subsequent peaks) and negative values indicate phase delays (later subsequent peaks) [32]. Period shifts were estimated as the difference in regression-derived cycle length, with regressions fit separately to the pre- and post-stimulation acrophase sequences. For multi-day entrainment experiments, phase angle relative to stimulation on the first day after stimulation ended was computed as the scheduled stimulation time minus the *Per2*-ELuc acrophase for that day (acrophase estimated from the post-stimulation regression), wrapped to (−12, 12] h. Positive values indicate peaks occurring before stimulation (phase-advanced relative to the stimulus), and negative values indicate peaks occurring after stimulation.

Phase response curves (PRCs) and period response curves (τRCs) were fit by modeling phase shifts or period shifts, respectively, as a smooth periodic function of stimulation CT using periodic harmonic regression on a Fourier basis with two harmonics. Stimulation events were assembled in long format with species (*Mus*, *Rhabdomys*), condition (experimental, control) and animal identifiers, stimulation CT (wrapped to [0, 24)), and the corresponding phase shift or period shift response. Species or condition comparisons used a single model with a group main effect and group-by-Fourier-term interactions, allowing the full periodic curve shape to differ by group while enforcing continuity and periodicity. Fitted mean curves were evaluated on a dense CT grid, and pointwise 95% confidence bands for the fitted mean were computed from the ordinary least squares coefficient covariance using the delta method.

Overall PRC shape differences between two groups (A and B, defined by species and/or experimental condition) were quantified with an animal-blocked permutation test on an integrated squared-distance metric computed across circadian time (CT):

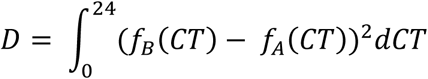

where f_A_(CT) and f_B_(CT) are the fitted mean PRCs for groups A and B, respectively [49,50]. *D* was approximated by trapezoidal integration over the CT grid. To account for repeated measures within animals, group labels were permuted at the animal level, the PRC model was refit for each permutation, and *D* was recomputed to form a null distribution (2000 permutations). The one-sided p-value was computed from the proportion of permutations with D_perm_ ≥ D_obs_ with a +1 correction in numerator and denominator.

PRC features were extracted from the fitted mean curves on the CT grid. Advance and delay areas were computed as the integrals of the positive and negative portions of the curve, respectively, using trapezoidal integration. Advance and delay centroid phases were computed as circular weighted means of CT using weights equal to the positive (advance) or negative (delay) mass, and resultant length was computed to summarize the concentration of the weighted distribution. Prespecified windowed delay metrics were computed for the early subjective day interval CT 2 to 8, used operationally as a “dead zone” window, by integrating or averaging the fitted curve or its negative component over the window.

Uncertainty and hypothesis tests for PRC features used animal-level resampling. Within-group confidence intervals were obtained by animal-resampled bootstrapping (2000 iterations), sampling animals with replacement, carrying forward all stimulations from sampled animals, refitting the PRC model each iteration, and re-extracting features from the refit mean curves. Between-group feature differences were tested using animal-blocked, two-sided permutation tests (2000 permutations) in which group labels were permuted at the animal level and feature differences were recomputed after refitting. Timing differences were wrapped to the shortest direction around the circle (for 24 h, to (−12, 12]) prior to inference. Holm multiple-comparison correction across extracted features was applied to the resulting per-feature p values. PRC and τRC fitting and resampling-based inference were implemented in Python (custom scripts; see Code Availability).

### Single cell calcium imaging

After viral expression of GCaMP6s, SCN slice cultures on membrane inserts were transferred to 35 mm glass-bottom wells in a 6-well plate (Cellvis) according to established methods [51]. The plate was sealed with optical adhesive film (Applied Biosystems) and placed in a light-tight environmental enclosure (Tokai Hit) maintained at 36°C under ambient air for time-lapse imaging. Imaging was performed on a Ti2-E inverted microscope (Nikon) equipped with a spinning disk confocal head (X-Light V3, Crest Optics), a motorized stage (Nikon), and a 20×/0.75 NA dry objective (Plan Apochromat Lambda D, Nikon). GCaMP6s fluorescence was excited with a 488 nm laser at 8% power and collected through a GFP filter set. Fluorescence was detected with a back-illuminated 10-megapixel sCMOS camera (Kinetix, Photometrics) using 1×1 binning. Single focal-plane images were acquired at 2394 × 2394 pixels (0.325 µm/pixel at the specimen; field size approximately 1.04 × 1.04 mm), 400 ms exposure, once every 15 min for 48 to 120 h, with the motorized stage sequentially visiting each slice and returning on the next 15 min cycle. Focus was maintained throughout recordings using Nikon Perfect Focus. Imaging parameters (laser power, camera gain, and acquisition settings) were held constant within an experiment.

### Single-cell calcium imaging analysis and phase mapping

Long-term single-cell calcium imaging movies were analyzed to estimate peak phase for tracked neuronal ROIs and to compare atlas-registered spatial phase organization across animals using binwise phase maps and cluster-based permutation inference.

ROIs were extracted and tracked using an extended, modified implementation of previously published code [52]. Briefly, candidate ROIs were detected independently in each frame using preprocessing designed to enhance soma-like features. Frames were filtered using difference-of-Gaussians filtering applied to the inverted image, and candidate ROI centroids were identified as local minima in the filtered image and ranked, with the highest-scoring candidates retained per frame. Candidate detections were then linked across time into trajectories based on spatial proximity, allowing brief dropouts. Trajectories were filtered to enforce plausibility, including requiring detections across a minimum fraction of frames, excluding trajectories that left the field of view, and excluding trajectories that contained implausibly large frame-to-frame jumps.

For each retained trajectory, a ROI fluorescence trace was extracted by sampling a local neighborhood around the tracked centroid at each frame. Trace extraction used a Gaussian-weighted intensity measure computed in a fixed local neighborhood centered at the ROI centroid, yielding an ROI-by-time matrix for downstream rhythm, phase, and visualization analyses.

ROIs were refined using automated morphology checks together with manual curation. Automated quality control used an ROI “cellness” score computed from a local image patch centered on each ROI. Soma-like morphology was evaluated using patch-based size and shape metrics (area and eccentricity) together with a center-versus-surround contrast metric (center disk compared against an annular neighborhood), using prespecified thresholds. ROIs were then reviewed alongside their morphology metrics, tracking stability, and extracted fluorescence trace with rhythm model overlays, and were excluded when they showed non-somatic morphology, unstable tracking, or traces dominated by artifacts.

Rhythm and phase were estimated from extracted ROI fluorescence traces using detrending, per-slice period discovery, and single-component Cosinor regression [53]. ROI traces were detrended by subtracting a running-median baseline computed with a 36 h window (144 frames at 15 min per frame). A single dominant period was estimated per slice from the FFT of the mean detrended signal computed across ROIs, selecting the peak-power frequency within a prespecified period range of 22 to 28 h. This slice-level period estimate was then used as a shared period for Cosinor fits to all ROIs within that slice. Rhythmic ROIs were defined by single-component Cosinor regression using the criteria p < 0.05 and r² ≥ 0.2, and peak phase was defined as the Cosinor acrophase expressed in hours and wrapped within one cycle.

Within each slice, phases were centered to yield relative phase by subtracting a reference phase (the circular mean across all rhythmic ROIs in the slice) using circular arithmetic. Relative phases were visualized as ROI point maps in which ROI centroids were colored by relative phase, and as gridded phase maps generated by interpolating the cosine and sine components of phase across a spatial grid and converting back to phase angle. ROIs outside a user-defined anatomical outline of the SCN were excluded from visualization and downstream spatial analyses.

To compare spatial organization across animals, each slice was registered into atlas space using landmark-based warping. Corresponding anatomical landmarks were placed in the slice image and in an SCN atlas image, a warp transform mapping dataset coordinates into atlas coordinates was computed, and the transform was applied to ROI centroid coordinates to generate atlas-registered ROI maps.

Atlas-registered ROIs were binned onto a regular 2D grid to construct binwise phase maps. For descriptive group maps, ROIs were pooled across animals after registration, and each bin value was the circular mean of ROI relative phases, with Gaussian spatial smoothing applied for visualization. For inference and phase-difference mapping, after converting relative phase to circadian time on a fixed 24 h cycle, binwise summaries were computed at the animal level first (per-animal circular mean per bin), then combined across animals to form group maps and group phase-difference maps, preventing animals with high ROI counts from dominating bin estimates. Bins were included only when they met prespecified minimum coverage criteria across animals and ROIs: bins required at least 5 animals per species for visualization and at least 9 animals per species for inference, with at least 10 cells per bin.

Spatially contiguous phase differences were identified using cluster-based permutation inference on the binned phase-difference map [54,55]. The per-bin statistic was the absolute circular phase difference between groups, computed on the 24 h circle from animal-level per-bin circular means. A cluster-forming threshold was set using a cluster-forming alpha criterion (α = 0.2), and suprathreshold bins were grouped into spatially contiguous clusters. Cluster mass was defined as the sum of per-bin statistics within each cluster. A null distribution for the maximum cluster mass was generated by randomizing group labels at the animal level and recomputing the maximum cluster mass for 10,000 permutations. Cluster-corrected p values were obtained by comparing observed cluster mass values against the permutation distribution of the maximum cluster mass, controlling familywise error across the full spatial map.

To summarize dorsomedial-to-ventrolaterally organization, atlas-registered ROI centroids were projected onto a dorsomedial-to-ventrolateral axis defined as a polyline in atlas space. Each ROI was projected to the nearest point along the polyline, and its position was expressed as normalized arc length along the axis. When left and right dorsomedial-to-ventrolateral axes were defined, each ROI was assigned to the nearer axis, and axes were oriented dorsomedial-to-ventrolateral for consistency across slices. The axis was divided into 10 equal-length segments in normalized arc-length coordinates, and per-animal circular mean relative phase was computed within each segment after converting phase to circadian time on a 24 h cycle for cross-animal comparisons.

### Statistical analysis

Statistical analyses were performed in GraphPad Prism, MATLAB, and Python. Unless otherwise noted, tests were two-tailed and statistical significance was assessed at α = 0.05. Where analyses involved repeated measurements within animals, permutation and resampling procedures were blocked at the animal level so that the animal was the unit of replication for inference. Data are summarized as mean ± SEM unless otherwise indicated, and exact p values are reported in the figure panels or legends where applicable.

GraphPad Prism was used for Welch’s unpaired t tests, mixed-effects models fit by restricted maximum likelihood (REML), and two-way repeated measures ANOVA with Geisser-Greenhouse correction. Post hoc multiple comparisons used Tukey’s or Sidak’s tests as indicated for the corresponding analyses.

MATLAB was used for circular statistics and regression-based analyses. Circular tests (Harrison-Kanji two-way circular ANOVA and Watson-Williams tests) used the Circular Statistics Toolbox [56], and linear regression and ANCOVA used built-in MATLAB functions.

Resampling-based inference and permutation tests were implemented using custom Python scripts. For PRC and τRC comparisons, overall curve-shape differences were tested with animal-blocked permutation tests on the integrated squared-difference distance between fitted mean curves, and PRC feature differences were tested with animal-blocked permutation tests with Holm correction across extracted features. Feature uncertainty intervals were estimated by animal-resampled bootstrapping. For atlas-space phase maps, spatial inference used cluster-based permutation testing with animal-level label randomization and max-cluster-mass control of familywise error.

### Code availability

Code used for the single-cell calcium rhythm analyses is available at https://github.com/jones-lab-tamu/neuron-tracker-suite/. Code used for phase response curve (PRC) comparison analyses is available at https://github.com/jones-lab-tamu/compare_prc/.

## ACKNOWLEDGEMENTS

This work was supported by National Institutes of Health Grant R35GM151020 (J.R.J.) and a research grant from the Whitehall Foundation (J.R.J.). We thank members of the Jones Lab and M. Tackenberg (University of Kentucky) for helpful discussions and feedback on the manuscript, R. Mallarino (Princeton University) and the Department of Biology vivarium staff (B. Earnest and M. Hooker) for invaluable support in establishing our striped mouse colony, D. Ferster (Actimetrics) for assistance with the LumiCycle optogenetics add-on, and M. Belle (University of Manchester) for guidance on single-cell calcium imaging.

