## Supplementary Figures for "Diurnality reconfigures circadian network dynamics in the suprachiasmatic nucleus"

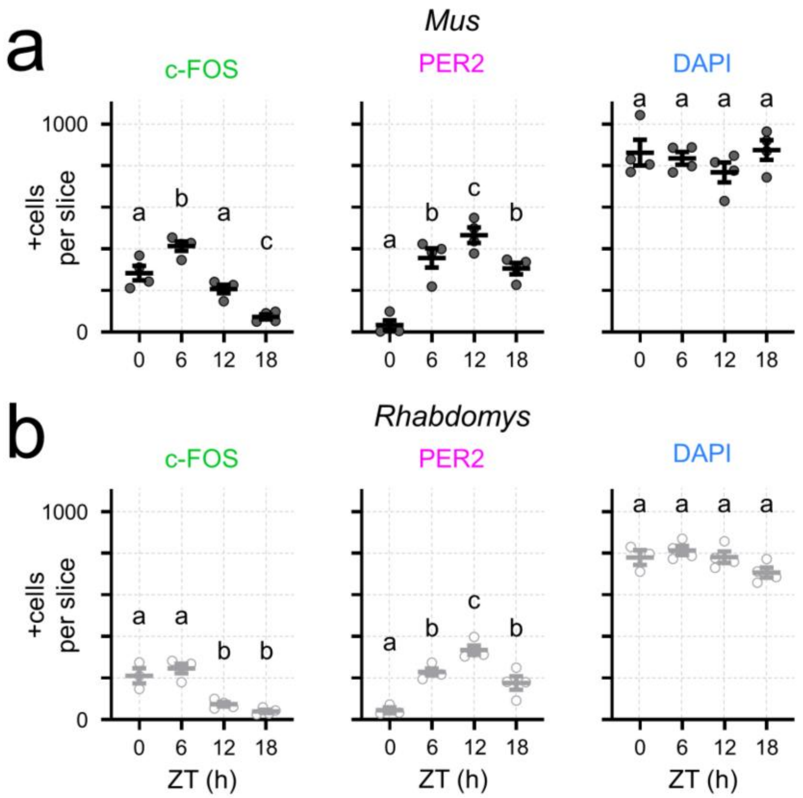


**Supplementary Figure 1. SCN c-FOS and PER2 cell-count rhythms show similar time-of-day ordering across species. a)** *Mus* SCN cell counts per slice across zeitgeber time (ZT 0, 6, 12, 18) for c-FOS$^+$, PER2$^+$, and DAPI$^+$ cells. Points represent individual animals (n = 4 per timepoint). One-way ANOVA: c-FOS, $F(3, 12) = 34.74$, $p < 0.0001$; PER2, $F(3, 12) = 28.18$, $p < 0.0001$; DAPI, $F(3, 12) = 0.9823$, $p = 0.4336$. Different letters indicate timepoints that differ by Tukey's post hoc tests ($p < 0.05$). **b**) *Rhabdomys* SCN cell counts per slice across zeitgeber time for c-FOS$^+$, PER2$^+$, and DAPI$^+$ cells. Points represent individual animals (ZT 0: n = 3; ZT 6, 12, 18: n = 4). One-way ANOVA: c-FOS, $F(3, 11) = 26.52$, $p < 0.0001$; PER2, $F(3, 11) = 24.07$, $p < 0.0001$; DAPI, $F(3, 11) = 3.040$, $p = 0.0746$. Different letters indicate timepoints that differ by Tukey's post hoc tests ($p < 0.05$). Across species, DAPI counts showed little evidence of time-of-day modulation, supporting that the rhythms in c-FOS and PER2 labeling primarily reflect changes in marker-positive counts rather than gross differences in the number of nuclei detected across timepoints.

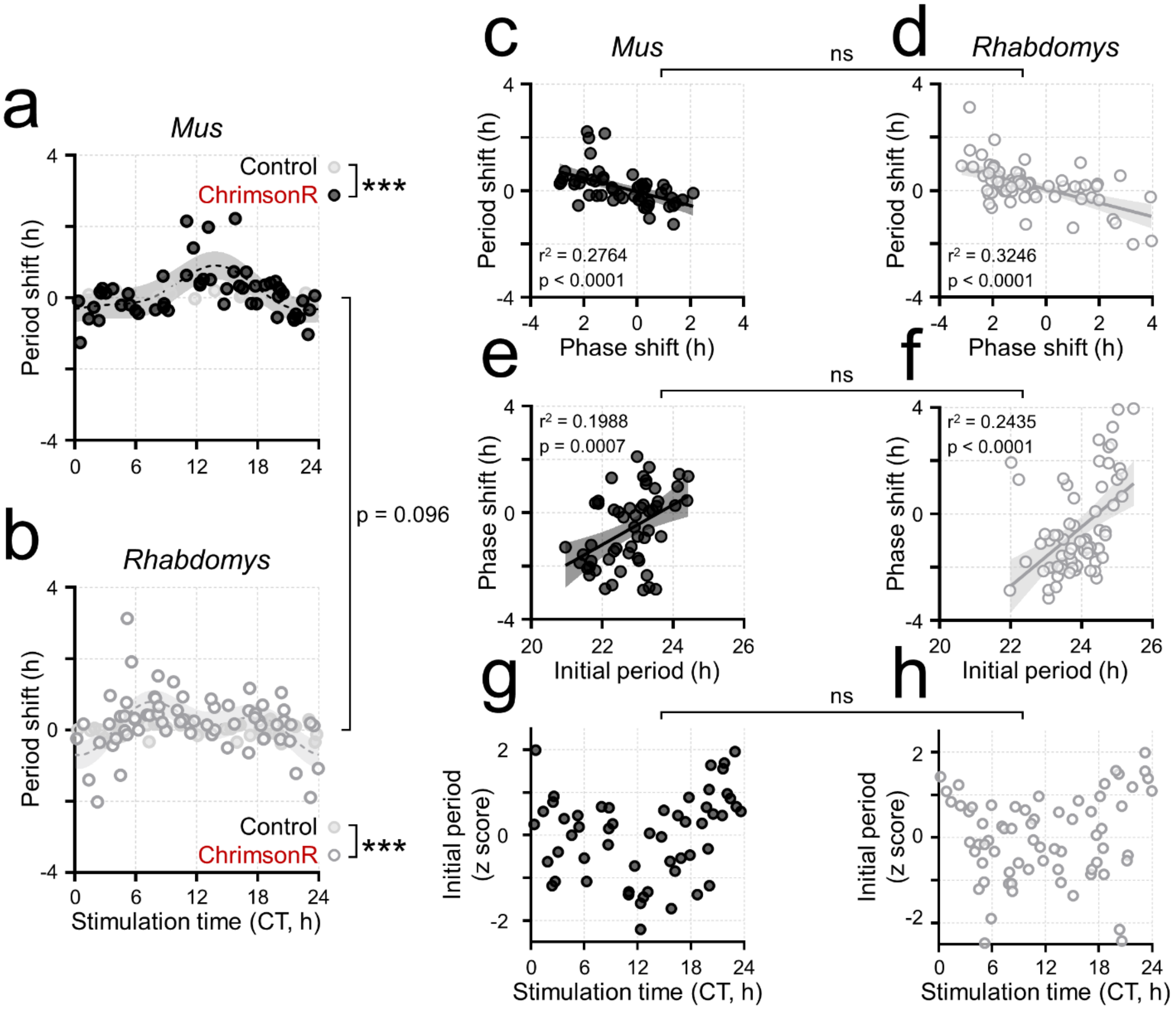


**Supplementary Figure 2. Period changes are unlikely to account for species differences in optogenetic phase resetting. a-b)** Period response curves for optogenetic stimulation of ChrimsonR-transduced SCN slices from nocturnal *Mus musculus* (**a**) and diurnal *Rhabdomys pumilio* (**b**). Points depict individual period shifts (in h; positive indicates period lengthening, negative indicates shortening) as a function of stimulus CT. Period shifts were quantified exactly as in Figure 2 (same regression-based estimator, inclusion rules, and stimulation dataset). ChrimsonR-transduced slices are shown as black filled circles (*Mus*) or gray open circles (*Rhabdomys*), and ChrimsonR-negative controls receiving identical light delivery and optical power are shown as light gray filled circles. Shaded bands indicate the 95% confidence interval of the fitted curve (periodic Fourier regression, k = 2). Ns are as in Figure 2. Curve shape differed between ChrimsonR and control slices within each species (animal-blocked permutation test on curve distance): *Mus* ChrimsonR vs. control, $D_{obs} = 4.536$, p = 0.0005; *Rhabdomys* ChrimsonR vs. control, $D_{obs} = 0.5643$, p = 0.0005. Curve shape did not differ between species (*Mus* vs. *Rhabdomys*), $D_{obs} = 5.1048$, p = 0.0970. **c-d)** Relationship between period shift and phase shift for optogenetic stimulations in *Mus* (**c**) and *Rhabdomys* (**d**). Points depict individual stimulations; lines depict simple linear regression with 95% confidence intervals. Period shifts covaried with phase shifts in both species (*Mus*: slope = -0.2578, $r^2 = 0.2764$, $F(1, 52) = 19.87$, $p < 0.0001$; *Rhabdomys*: slope = -0.2579, $r^2 = 0.3246$, $F(1, 65) = 31.24$, p

$< 0.0001$). The species by covariate interaction was not significant (ANCOVA; $F = 3.82 \times 10^{-6}$, $p = 0.9984$). **e-f)** Relationship between phase shift and initial pre-stimulation period for optogenetic stimulations in *Mus* (**e**) and *Rhabdomys* (**f**). Points depict individual stimulations; lines depict simple linear regression with 95% confidence intervals. Phase shifts covaried with initial period in both species (*Mus*: slope = 0.7525, $r^2 = 0.1988$, $F(1, 52) = 12.9$, $p = 0.0007$; *Rhabdomys*: slope = 1.112, $r^2 = 0.2435$, $F(1, 65) = 20.93$, $p < 0.0001$). The species by covariate interaction was not significant (ANCOVA; $F = 1.213$, $p = 0.2730$). **g-h)** Initial per-stimulation period (z-scored within species) versus stimulation time (CT) for optogenetic stimulations in *Mus* (**g**) and *Rhabdomys* (**h**). Points depict individual stimulations. Species did not differ in the stimulation-time dependence of the initial period (two-sided animal-blocked permutation test on the distance between $k = 2$ periodic Fourier fits; 2000 successful permutations; $D_{obs} = 9.808$, $p = 0.1499$).

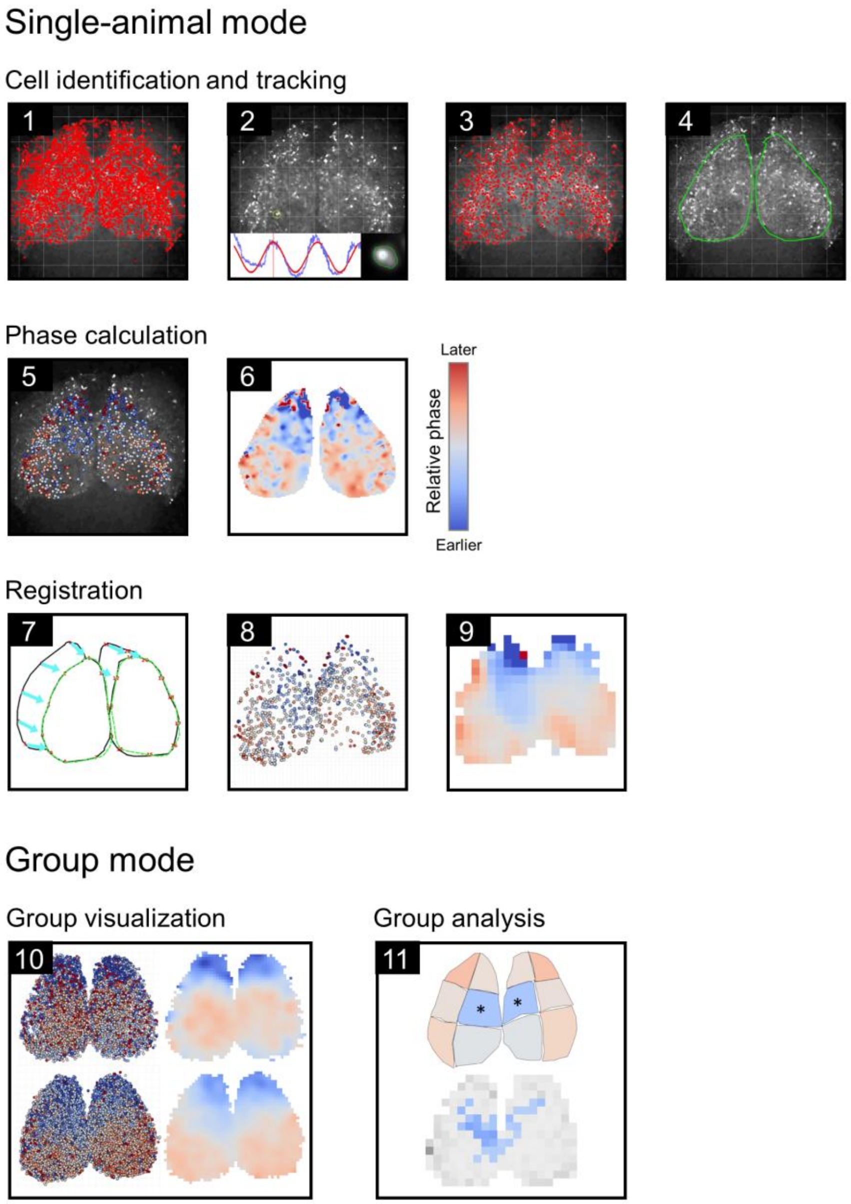


**Supplementary Figure 3. Application workflow for per-animal phase mapping, atlas registration, and group-level visualization and analysis.** 1) Candidate ROIs are detected and tracked from imaging movies. 2-3) ROIs are refined and quality-controlled using per-ROI metrics and visual inspection, including ROI “cellness” and fluorescence traces with model overlays for curation and rhythm extraction. 4) Anatomical boundaries are manually outlined on the background image to define regions for downstream analyses. 5-6) Peak phase is estimated per ROI from fluorescence traces and converted to relative phase by subtracting a circular-mean reference computed from all rhythmic ROIs or from a user-defined reference subregion. Relative phases are visualized as ROI point maps and as gridded phase maps. 7) Atlas registration is performed by placing corresponding landmarks and computing a warp from dataset coordinates into atlas space, saving the transform for reuse. 8) The warp is applied to ROI coordinates and associated phase values to generate atlas-registered ROI maps. 9) Registered ROIs can be binned to generate binwise heatmaps for visualization and group comparisons.

10) For cross-animal comparison, relative phases are converted to circadian time on a fixed 24 h cycle, then displayed as pooled atlas-registered ROIs and binned heatmaps. 11) Group analysis computes region-level summary statistics and performs cluster-based permutation inference to identify spatially-clustered differences in binned phase-difference maps.

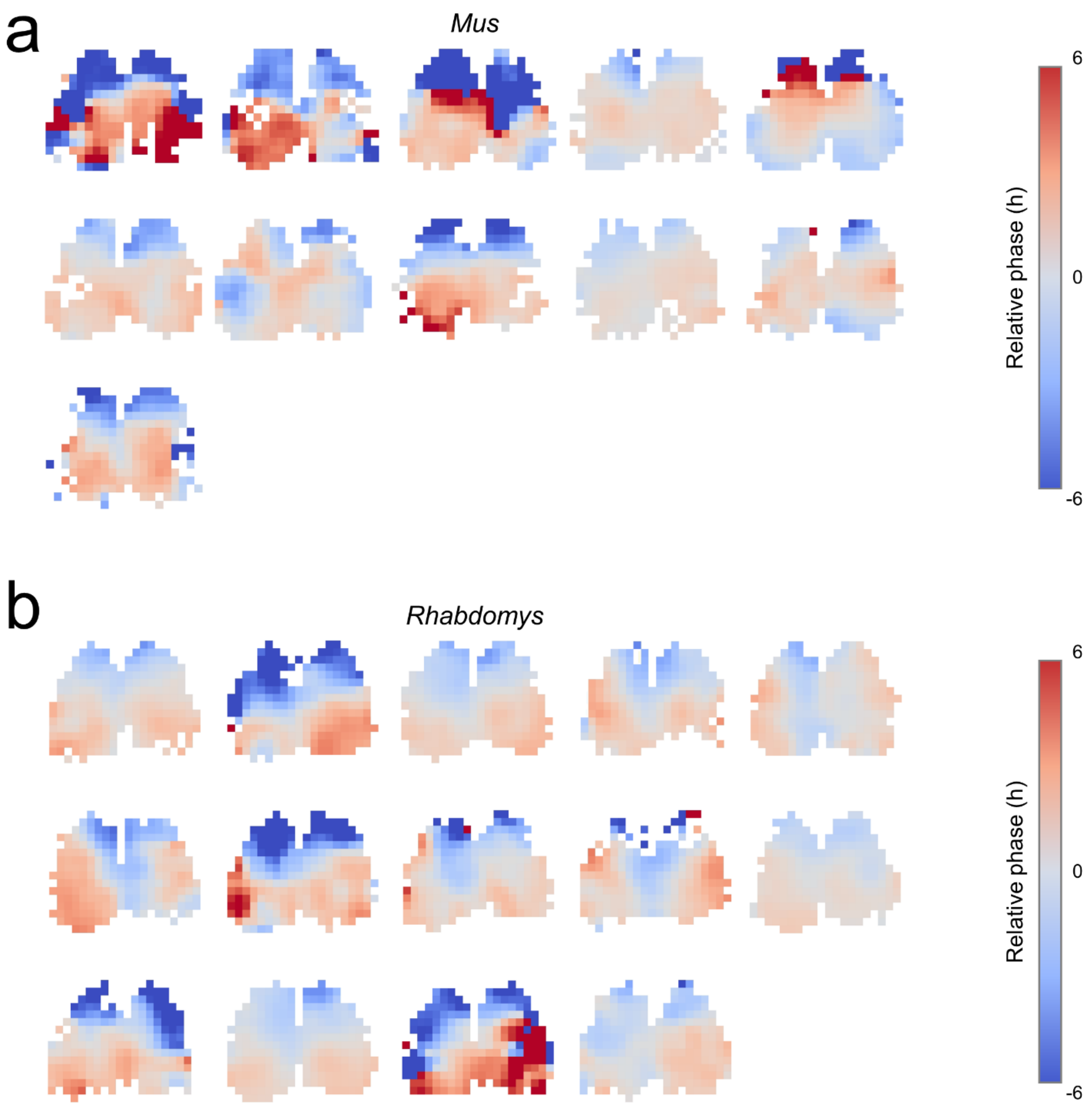


**Supplementary Figure 4. Individual-animal relative phase maps underlying group-averaged SCN phase maps. a-b)** Atlas-registered, binned relative phase maps for each SCN contributing to the group maps in Figure 3f (*Mus*: n = 11 animals; *Rhabdomys*: n = 14 animals). ROI-level relative phases were aggregated within spatial bins and displayed as 20 x 20 heatmaps to visualize across-animal variability in spatial phase organization.
